# In-Flights of Outbreak Populations of Mountain Pine Beetle Alter the Local Genetic Structure of Established Populations a Decade After Range Expansion

**DOI:** 10.1101/2024.04.02.586325

**Authors:** Kirsten M. Thompson, Dezene P. W. Huber, Felix A. H. Sperling, Brent W. Murray

## Abstract

Mountain pine beetles began to appear at epidemic levels in Alberta, Canada, in 2006, following six years of extensive outbreaks in neighboring British Columbia. We assessed the effect of genetic MPB in-flights from the peak of the outbreak on the genetic structure of established populations of MPB and the change over time in novel regions colonized by these inflights. We used five locations sampled during the peak of the outbreak (2005/2007) and re-sampled in 2016. We performed a ddRADseq protocol to generate a SNP dataset via single-end Illumina sequencing. We detected a northern and southern genetic cluster in both sampling sets (2005/2007 and 2016) and a demographic shift in cluster assignment after ∼10 generations from south to north in two of the sites in the path of the northern outbreak. Fst values were significantly different between most sites in the same years and between the same sites at different years, with some exceptions for northern sites established by inflights. Overall, sites in the spreading path of the MPB outbreak have taken on the genetic structure of the contiguous northern outbreak except for an isolated site in Golden, BC, and in Mount Robson Provincial Park where populations are admixed between north and south. Our results suggest that range expansion during insect outbreaks can alter the genetic structure of established populations and lead to interbreeding between populations.

## Introduction

Global ecosystems are experiencing dramatic changes in climate regime, leading to species from multiple taxa undergoing range shifts or range expansions (Parmesan, 2006; Clements & DiTommaso, 2012). In some cases, these climatic changes lead to detrimental effects and species loss due to habitat exclusion, but in other cases species benefit from climatic changes by experiencing population growth and increased movement (Ciosi *et al*., 2008; Mona *et al*., 2014). Model-based genetic studies suggest that range expansion causes established population structure to change over time, with changes in patch occupation, gene flow between populations, and allele surfing contributing to alterations in genetic character (Klopfstein *et al*., 2006; Mayrand *et al*., 2019). Populations on the leading edge of a range expansion may also lose structure as they carry only a subset of the genetic diversity of the species, while at the same time establishing the foundational population throughout the new habitat range (Ibrahim *et al*., 1996).

The effects of range expansions and invasion often focus on phenotypic changes or population demographic shifts to document the changes a species undergoes after migration to a new area (Young *et al*., 2017). Short-term genetic consequences of range expansions are addressed less frequently in the literature. Instead, many studies focus on historical genetic changes comparing geological time scales or movement away from glacial refugia (Hellberg *et al*., 2001; Roberts & Hamann, 2015; Hagen *et al*., 2015). The studies that do document genetic impacts over relatively short periods of time (several years to decades) focus on population structure changes within established species ranges, relying on archived samples and citizen collections with imprecise location data, as seen with studies of red deer (Nussey *et al*., 2005) and bobcats (Carroll *et al*., 2019). Other genetic research on recently expanded species ranges utilises contemporary samples from the current population only to assess the establishment of current structure, as seen in the study of invasive crabs (Herborg *et al*., 2007), sparrows (Liebl *et al*., 2013), geckos (Short & Petren, 2011), and wasp spiders (Krehenwinkel *et al*., 2016). Rarely are the short-term temporal ramifications of species range expansion documented among years across repeated sites, as was the case in a study of brown bears that demonstrated the rapid genetic changes possible during range alterations (Hagen *et al*., 2015). In many cases, these genetic studies of range expansion focus on macrofauna that have relatively small numbers of offspring, or somewhat limited dispersal capabilities. Irruptive insects that produce high numbers of individuals that can disperse easily by air are not widely represented in the literature.

Mountain pine beetle, *Dendroctonus ponderosae* Hopkins (Coleoptera: Curculionidae) (MPB)) is one such irruptive insect. MPB is a tree killing sub-cortical pine bark beetle, responsible for one of the most widespread and damaging insect outbreaks in recent history (Raffa *et al*., 2008; Six & Bracewell, 2015). Over 18.3 million hectares of pine forests were affected by MPB in the wake of this outbreak in British Columbia from 2000–2012 (McKee *et al*., 2015). This native beetle is found in pine species in the southwestern USA to the Black Hills of South Dakota, most of British Columbia (BC), and recently expanded into Alberta. In western Canada, mountain pine beetle has been a common part of the natural disturbance regime of pine forests with recorded outbreaks of differing size recorded throughout the 20^th^ century, and dendrochronologically reconstructed outbreaks dated prior to European settlement (Wood & Unger, 1996; Hrinkevich & Lewis, 2011).

Warmer temperatures linked to climate change prevented MPB winter brood die-off and increased breeding success in the early 1990s and into the 2000s (Safranyik & Linton, 1998; Carroll *et al*., 2006). Fire suppression and lack of human interference within established pine stands in favor of focus on more lucrative softwood species created a contiguous food source for spot outbreaks to grow and coalesce together (Whitehead *et al*., 2001). Historically, the mountainous terrain of the eastern portion of BC and the Rocky Mountains acted as range barriers to prevent excessive movement of MPB into Alberta, except in the areas around Banff National Park, the Crowsnest Pass, and Cypress Hills (Powell, 1961; Langor, 1989). MPB populations within the southern portion of BC were typically isolated from one another by the steep mountain terrain and patchiness of pine habitat interspersed with interior grasslands.

Mountain pine beetles began to appear in large numbers in Alberta in 2006 (Bartell, 2008), crossing over the Rocky Mountains assisted by strong convective winds (Jackson *et al*., 2008). These initial inflights of beetles were concentrated near Canmore in the south and Grande Prairie in the north but have since spread eastward as far as Lac La Biche (Bartell, 2008; Cullingham *et al*., 2011; Pokorny, 2021). A decade after the initial colonization of these regions during the peak of the outbreak, beetles are still present and continuing to attack surviving pine. Some areas appear to be experiencing epidemic-level attacks while others appear to be collapsing down to endemic levels.

Several genetic studies addressed the population structure of MPB within western Canada using a variety of marker systems. One of the first within BC used microsatellite markers to investigate the MPB outbreak throughout its original range and parts of the expanded range (Bartell, 2008). Populations in the north were found to be more homogenous, while southern populations were more structured, though although the authors did not identify a landscape barrier leading to this division (Samarasekera *et al*., 2012). A similar study using single nucleotide polymorphisms (SNPs) confirmed the differentiation between the northern and southern geographic clusters, also finding that the Robson Valley and nearby Jasper National Park had greater genetic diversity in an intermediate area between the north and south (Samarasekera *et al*., 2012; Janes *et al*., 2014; Batista *et al*., 2016). These studies combined beetles from multiple years, but also were conducted during or within several years of the initial MPB range expansion in the mid-2000s. In all cases, the researchers did not have the opportunity to re-sample regions and compare MPB genetic structure between age cohorts.

It is unknown if the populations of MPB in regions that received a high level of migrant beetles, like Smithers or Canmore, retained their original endemic population structure 10 years after the initial in-flights, or if they have taken on the character of the northerly MPB population that expanded into the north of BC and into Alberta. It is also unknown how locations with novel colonization, like Grand Prairie, have changed since the first in-flights. In both cases, beetles will have completed approximately nine to eleven generations at each location, as MPB is known to re-attack relatively close (<3 km) to their original host tree if conditions are favourable (Evenden *et al*., 1943) and the correct aggregation pheromone cues are received (Evenden *et al*., 2014).

Our study took advantage of beetles archived by research teams during the first wave of expansion of the BC outbreak in 2005 and 2007 and compares those specimens to populations active in 2016. Our objective was to document the short-term temporal genetic structure changes that occur in the aftermath of an insect range expansion with a high degree of immigration into novel territory and into areas with established populations. Temporal genetic sampling methods have been recommended for use in assessing genetic structure post-range expansion (Short & Petren, 2011; Hagen *et al*., 2015), though their use in insects at this time has largely been restricted to agricultural pests responding to abiotic inputs (Pélissié *et al*., 2018). We hypothesized that samples from the first time point would have greater genetic structure than those sampled ten years on, but that samples from established populations would retain, in part, their original structure due to dispersal limitations. We also anticipated that admixture would increase over time due to gene flow between sampling locations (Hagen *et al*., 2015).

## Methods

### Sampling and Extraction of Genomic DNA

Field sampling was conducted in the summer of 2016. Adult and teneral (un-hardened) beetles were collected from trees in British Columbia and Alberta. Sampling was informed by aerial survey data provided by the BC Ministry of Forests, Lands and Natural Resource Operations (https://catalogue.data.gov.bc.ca/, 2022) and field survey data from Alberta Agriculture and Forestry (C. Whitehouse, personal communication, June 2016). Infested trees were identified in the field by observing yellowing foliage and the presence of pitch tubes accompanied by frass on the bole. Bark was peeled using a draw knife to expose pupal chambers. A minimum of 40 adult or teneral beetles were collected per site, with individuals selected from separate galleries to prevent comparison of full siblings. Samples collected from 2005 and 2007 used the cardinal direction collection method, where beetles were selected from four points around the bole of the tree to prevent gallery overlap. In both cases, a maximum of four per tree were retained when possible. Beetles collected in 2016 were immediately placed in 95% ethanol upon removal from the tree in the field, transferred to a lab, and then stored at -20℃ prior to DNA extraction. All the 2005 and 2007 specimens from Alberta and BC used in this study followed the same method of collection and ethanol preservation as above. Samples from 2007 were collected as second instar larva and lyophilized for room-temperature, shelf-stable storage prior to DNA extraction.

Genomic DNA was isolated from all MPB specimens collected in 2016, and those collected in 2007 using Qiagen Dneasy Blood & Tissue spin column and 96-well plate kits (Germantown, Maryland, USA) following manufacturer’s instructions, with the addition of an overnight tissue lysis step incubated at 56℃ and the use of an optional RNA removal step using 2µL of RNaseA with a 15-minute incubation. DNA product was eluted into DNase-free water for ease of later sequencing. DNA from the samples collected in 2005 was previously extracted using a phenol-chloroform method and stored at –80℃ (Samarasekera *et al*., 2012). Final concentration was quantified using both a NanoDrop™ 1000 Spectrophotometer and a first-generation Qubit fluorometer with the Quant-iT dsDNA BR Assay Kit.

### Sequencing and Data Cleaning

Double-digest restriction-site associated DNA sequencing (ddRADseq), a type of genotyping-by-sequencing, was used to generate the SNP markers for this project. Genomic library preparation and sequencing was completed by the Molecular Biology Services Unit (MBSU) of University of Alberta. DNA Library preparation followed the MBSU specifications for MPB sampling (Peterson *et al*., 2012; Campbell *et al*., 2017). Total purified DNA input per sample was 80ng. A non-methylation sensitive two-enzyme system (*Pst* I and *Msp* I) was used to fragment whole genomic DNA. Samples were sequenced on a single lane of an Illumina NextSeq 500 at the MBSU to generate single-end 75 bp reads.

Initial sequence read demultiplexing and mapping was performed on Compute Canada’s Graham cluster (Toronto, Ontario, Canada). Reads were demultiplexed and quality checked using the default pipeline in STACKS 2.0b (Rochette *et al*., 2019). Cutadapt (version 1.9.1) was used to trim the *Pst* I restriction site on the 5’-end of each read (Martin, 2011). Reads were also trimmed to a length of 67 bp. Processed reads were mapped to the female draft MPB reference genome (Keeling *et al*., 2013) using BWA-MEM (Burrows-Wheeler Aligner Maximal Exact Match – version 0.7.17) (Li & Durbin, 2009). Quality of the alignments was assessed using SAMtools (version 1.9) (Li *et al*., 2009b).

Single nucleotide polymorphism variants were called by the ref_map.pl perl wrapper provided with STACKS 2.0b with a minimum minor allele frequency of 0.05. We also used the r80 principle where any retained locus must appear in at least 80% of all individuals (Paris *et al*., 2017; Rochette *et al*., 2019). VCF files generated by ref_map.pl were filtered again using VCFtools (version 0.1.14) for a minor allele frequency of 0.05, a minimum quality score of 30, and minimum read depth of 10 (Danecek *et al*., 2011). To mitigate the confounding signal of sex-linked markers, the dataset was loaded into R (version 3.6.3) and SNP loci were filtered using the PCA based method in (Trevoy *et al*., 2019) by removing loci associated with high PC 1 values (R Core Team, 2018). We filtered for linkage disequilibrium (R^2^ > 0.5) using the R package dartR (Abdellaoui *et al*., 2013; Gruber *et al*., 2018) retaining only one random SNP from each linkage group to prevent excessive clustering during population structure analysis. Individuals were rechecked for quality after linkage filtering and any remaining poor-quality individuals (more than 20% data missing) were removed.

### Assessment of Population Structure

SNP data were first visualized using the adegenet package (version 2.1.2) in R (Jombart, 2008; Jombart *et al*., 2010). A combination of PCA and DAPC was used to assess similarity between the two time periods and between the individual locations. PCA was performed without assigning populations to specific groups based on collection year. DAPC maximizes the variation between groups and uses a priori population assignment. Individuals were assigned to populations corresponding to their location of origin. Cross validation calculations were performed following the defaults contained in the DAPC vignette to assess the number of PCs to retain and the DF to retain for final visualization (Jombart *et al*., 2010).

We used STRUCTURE (version 2.3.4) to explore population structure among all of our populations (Pritchard *et al*., 2000; Falush *et al*., 2003, 2007; Hubisz *et al*., 2009). For this dataset, we used the admixture model and did not specify location information or group individuals into populations. We performed an initial STRUCTURE run with 100,000 Markov chain Monte Carlo generations and a burn-in period of 50,000. We then increased our STRUCTURE run to 1 million Markov chain Monte Carlo generations and a burn-in period of 250,000 and tested for values of K between 1 and 10 with 10 replicates of each K value.

STRUCTURE results were visualized using CLUMPAK (Kopelman *et al*., 2015) to create an average for each value of K. Optimal K was determined by comparing the results of the ΔK Evanno method (Evanno *et al*., 2005) between likelihood of K values and the mean estimated natural logarithm for the K value probability (Ln(PrK))(Pritchard *et al*., 2000). Both ΔK and Ln(PrK) were compared using STRUCTUREHARVESTER (Earl & VonHoldt, 2012). A hierarchical AMOVA was conducted in Arlequin (version 3.5.2.2) using pairwise Fst comparisons on the 10 sites, among the two collection dates, among sites within the collection dates, and among individuals within populations, and within individuals, set at a 0.05 significance level with 1000 permutations (Excoffier & Lischer, 2010). Isolation by distance was tested using a Mantel test in dartR (version 1.1.11) with a Pearson’s product-moment correlation and a Fst/1-Fst vs. the log distance in meters based on the Mercator projection (Gruber *et al*., 2018). Summary statistics of the cleaned datasets were conducted using GenoDive (version 3.05) to assess observed heterozygosity (Ho), heterozygosity within populations (Hs), and inbreeding coefficients (*Gis*) (Meirmans, 2020).

## Results

We retained 4,899 genomic SNPs and 175 individuals within the dataset after quality filtering. Pairwise Fst values ranged from 0 to ∼0.14 and were largely significantly different population-to-population, regardless of collection year (Table 2.2 and Table 2.3). The exceptions to this were the two Grand Prairie timepoints (2007 and 2016) which were non-significant, the initial Canmore and Golden site sampling (2007 and 2005 respectively), and the initial 2007 Grand Prairie sampling and the 2016 Canmore sampling. Observed heterozygosity (Ho) was comparable on all sites (0.293 – 0.265) with no major trends connected to sampling date. The highest Ho was detected at the Golden site from 2007. Inbreeding coefficient values ranged from 0.093 to -0.036 and were highest at the 2016 collection at Canmore (Table 2.1).

Both ΔK (Figure 2.3) and Ln(PrK) metrics from our STRUCTURE analysis support an optimal K of 2 at both time points, indicating two detectable population clusters on the landscape (Figure 2.2). Canmore showed a southern affinity during the first sampling and a mixed northern and south affinity during the second sampling. Smithers at the time of the first sampling showed a more mixed population and in the 2016 sampling grouped completely north. In contrast, Robson moved from a blend of individuals assorting fully to south and north, to most individuals having probability of cluster membership divided evenly between north and south (this admixed signature is addressed more fully in Chapter 3). Golden remained southern in nature at both sampling times, though there was a low-level increase in probability of northern assignment.

Finally, Grande Prairie maintained a northern population assignment at both sampling times. An additional cluster (K = 3) is not supported by the data and additional K values beyond 3 are also not supported. Clusters were also assessed using the find.clusters algorithm in adegenet which relies on Bayesian information criterion (BIC) to infer optimal populations structures. K = 2 was also supported in this analysis. Isolation by distance analysis of sites grouped by both year and taken together showed no pattern of geographic population differences by distance, r^2^ = 0.2189 (Figure 2.5).

DAPC and PCA analysis echoed the results of the STRUCTURE analysis with association of populations with collection year, but more association with geographic clustering. In the case of both STRUCTURE and DAPC (80 PCs retained), sites that were in the expanding path of the BC outbreak took on a more northerly signature (ellipses overlapping), with Mount Robson Provincial Park taking an intermediate position and Golden remaining distinct (Figure 2.4C). PCA investigation of both time points taken together showed roughly the same geographic clustering reflected in the STRUCTURE analysis, with PC 1 explaining 9.68% of the variation found in the SNP dataset and PC 2 explaining 4.5% of the remaining variation. PC 1 is likely linked to geographic location, while PC 2 is most likely linked to variation between individuals (Figure 2.4A) (Shegelski *et al*., 2021). Analysis of molecular variance (AMOVA) of 4733 loci was used to test the differences between sites and to look for similarity based on year or location (Table 2.4). Our analysis found that there was considerable variation within populations and between individuals. Grouping populations by year explained the least amount of variation. All Fstatistics sampled (Fis, Fsc, Fct, Fit) were significant (p < 0.0001).

## Discussion

Our study assessed the changes in population structure among MPB populations sampled during the height of the British Columbian outbreak of the early 2000s and those sampled approximately ten years after expansion and of the initial population to the north and east. We explored the possibility that beetles would retain their original population signature in the path of an expanding outbreak due to dispersal limitations based on their flight capacity and proximity to attractive viable host trees. We found instead that populations in the path of the outbreak (Canmore in the east to Alberta and Smithers in the north) transitioned from more southerly or mixed assignments to more northernly assignments and took on the characteristics of the large spreading BC outbreak. However, beetles in all collection years across all sites still had a distinguishable north-south population divide, with two detectable genetic clusters on the landscape that assorted geographically, even though there was no discernible isolation-by-distance detected. It is likely that the smaller number of total individuals in this study and the greater number of sites that assort to the north obscured any pattern of isolation by distance. This echoes previous research on MPB in western Canada that also has shown a geographic north/south divide between populations (Samarasekera *et al*., 2012; Janes *et al*., 2014; Batista *et al*., 2016; Shegelski *et al*., 2021).

The 2016 samples from Mount Robson Provincial Park population had an intermediate split of populations assigned between north and south for all individuals. Mount Robson Provincial Park is located close to the area that was identified by Janes *et al*. (2014) as region of higher genetic admixture within BC and as an area of genetic mixing by Samarasekera *et al*. (2012). The original sampling of Robson in 2005 produced more individuals that assorted completely north and completely south (a mixed population) as opposed to these more admixed individuals collected in 2016 (Figure 2.2). This indicates that the Robson area in 2005 contained beetles from both landscape clusters that have likely since interbred to produce a blended population. It is important to note that STRUCTURE analysis did not identify a distinct population in this area as a separate cluster, nor did the find.clusters algorithm with this dataset.

AMOVA analysis of the SNP dataset indicated that while MPB individuals have high genetic variability among each other, both populations and collection-time based groupings produced small but significant differences. The recent expansion of the MPB outbreak, combined with the presence of two population clusters and mixed populations within the dataset likely reduced the size of the difference between the two time points, but it is important to recognize that there are changes in the geographic population structure between 2005/2007 and 2016.

There are significant site-to-site pairwise Fst differences over time in almost all sites sampled in 2005/2007 and 2016, likely reflecting a history of differentiation due geographic isolation with limited long-distance dispersal between populations prior to the large MPB outbreak in BC during the mid-2000s.

The mid-2000s outbreak moved both north and east, leaving southern BC outside of the path of most dispersing MPB in the north of the province. In addition to the Golden population’s isolation in a high, remote mountain valley in the south of BC, the outbreak’s movement away from the southern locations following prevailing winds likely contributed to the maintenance of the distinct southern structure detected at Golden (Figure 2.4 B and C). The two Grande Prairie MPB collections in the north of Alberta did not differ by pairwise Fst, likely due to the relatively recent establishment of the population and the fact that all MPB populations in the expanded range area come from the northern BC epidemic. Approximately ten generations in one location in a region without repeated population replenishment from other MPB sources is likely insufficient to cause a significant change in the original genetic structure.

In most of British Columbia, MPB populations have now returned to endemic levels due to lack of food resources and extremes of temperature in the expanded range in the north of the province. Alberta has seen similar declines due to very vigorous control efforts, and a series of non-ideal climate conditions including excessively cold winters in 2019 and 2020, and an unseasonably cool and wet summer in 2019 (M. Undershultz, personal communication, April 22, 2022). The removal of the host pine by several consecutive summers of intense wildfire in both provinces is also likely driving beetle numbers down, returning both areas to an endemic level of infestation (Tan *et al*., 2019; Daniels *et al*., 2020).

MPB populations in northern Alberta, specifically in the Grand Prairie area and eastward, are considered locally invasive as there has never been a recorded instance of MPB colonization prior to 2006 (Burke, 2016). This is despite the fact that the Canmore population in southern Alberta is connected to the well-established Banff populations that have been documented since the 1940s (Powell, 1961; Cooke & Carroll, 2017). MPB in Canmore has taken on the more northernly genetic signature suggesting that gene flow toward the park is significant enough to prevent new local adaptations from establishing at this time (Allendorf *et al*., 2013). In addition, the presence of northernly beetles in Canmore may indicate that the northern population is beginning to destabilize existing genetic structure in the Banff National Park area.

Despite the shorter geographic distance between Robson, located between north and south clusters, and Canmore compared to Grand Prairie and Canmore, STRUCTURE grouped most MPB individuals in Canmore in the south with Grande Prairie in the north which may indicate either long-distance dispersal events from the colonizing MPB populations in Alberta or local transportation of infested wood into the area. Transportation of firewood is discouraged specifically to prevent movement of pest insects (Government of Alberta, 2008), but Banff National Park and the surrounding towns accommodate a large number of tourists with demands for camp wood that may be sourced from infested areas in the north of BC or Alberta (Cheng, 1980). There is also the continued possibility of long-distance dispersal throughout Alberta naturally, as prevailing winds in all seasons move west to east through the north of the province, and down towards Canmore in the summer (Government of Alberta, 2014).

Though MPB outbreaks within the expanded range in central Alberta are in a state of flux, our study indicates that epidemic groups of beetles can establish a genetically homogenous population on a landscape even when separated by great distances. In the case of Alberta, the first colonizing wave of MPB from the northern BC outbreak has left a strong signature on the landscape. We also have found that there can be shifts in population assignment between MPB populations over short periods of time, meaning that care should be taken when combining sample cohorts from different years, particularly for irruptive species like MPB.

We found that regional genetic structure can be lost or altered in the face of epidemic level in-flights of beetles. Most of the beetles found in the northern parts of both BC and Alberta assort to one homogenous population. For this reason, the development of population-specific control methods should not be used over established methods of control as MPB populations on the landscape are likely not yet displaying meaningful behavioural differences site to site.

Traditional MPB management methods tailored to regional differences in stand compositions and site characteristics including: spot eradication through fall and burn, sanitation cuts, controlled burns, and other forms of host denial (Fettig *et al*., 2014) are all likely to be effective on current outbreaks.

The genetic characteristics of sites like Grande Prairie and Canmore are also important to document as they provide context for the character of MPB populations that are likely to remain in Alberta. To our knowledge, this is the only study to date that has compared the population genomics of an irruptive beetle pest by assessing genetic changes by time cohort. As repeated collections are now encouraged to track changes in genetic demography (Hagen *et al*., 2015), our data support the separation of collection cohort to better understand how irruptive pest movements are influencing population structure.

**Figure Error! No text of specified style in document..1:**
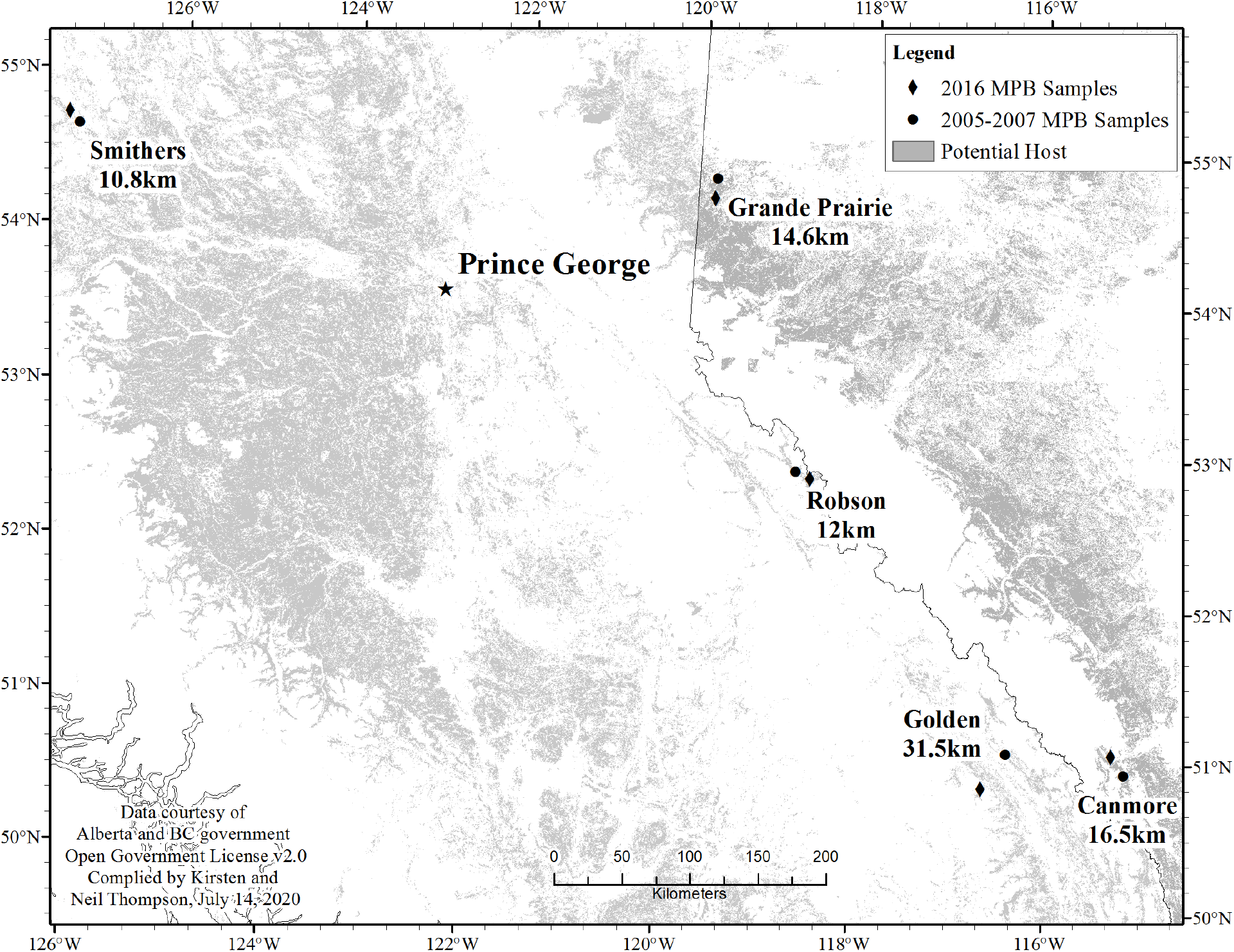
Map of MPB collection sites sampled in both 2005/2007 and 2016 with the distances between site locations.

**Figure Error! No text of specified style in document..2:**
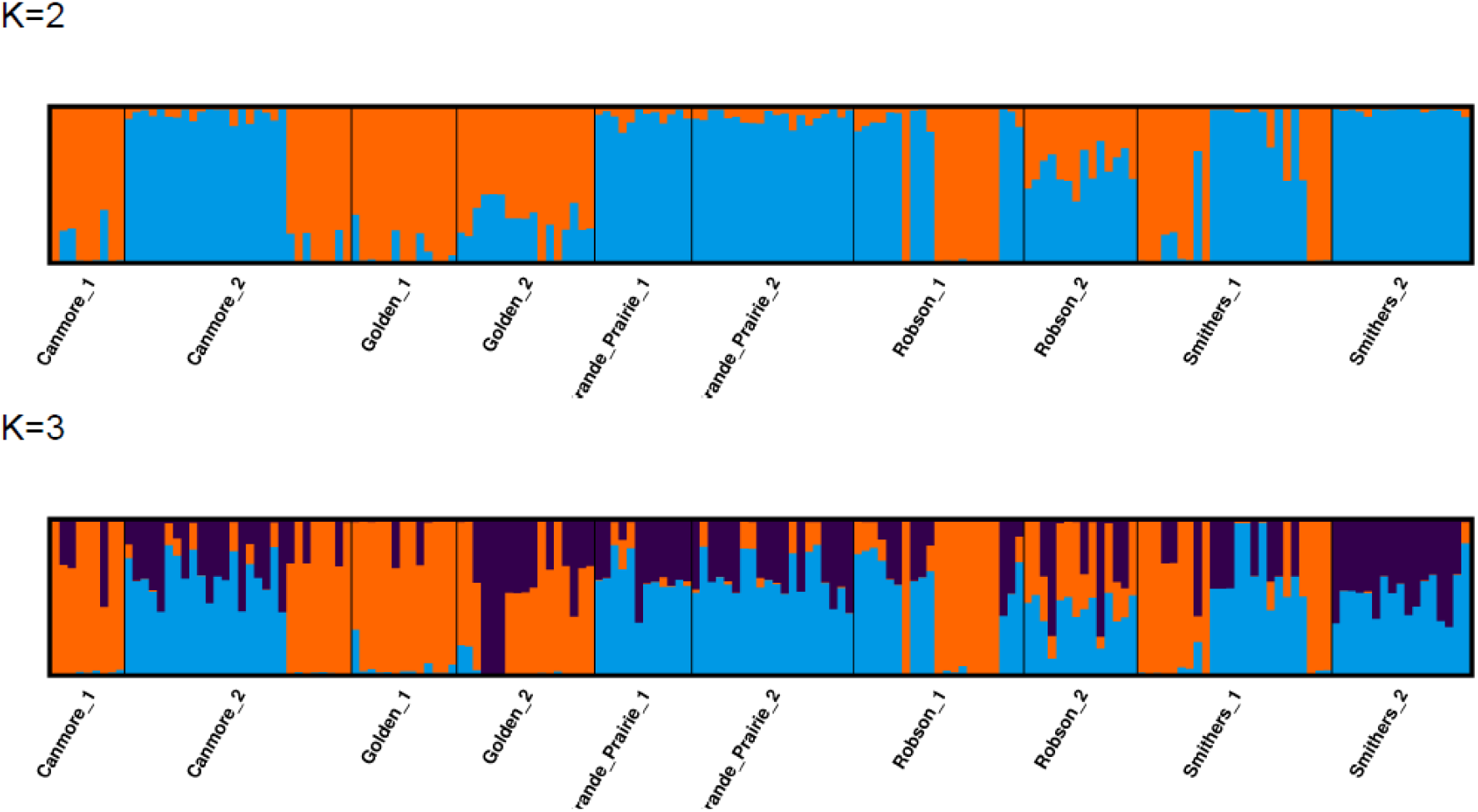
STRUCTURE plots with cluster membership per individual MPB (n=175) averaged by CLUMPAK. Sites visually divided by a black bar and, paired together by year, and analyzed at 4899 loci. Individuals are represented by a partitioned vertical bar with cluster membership colour coded by K. The proportion of colour within the bar represents the probability of the individual’s assignment to each cluster. For K = 2, orange denotes a southern cluster while blue denotes a northern cluster.

**Figure Error! No text of specified style in document..3:**
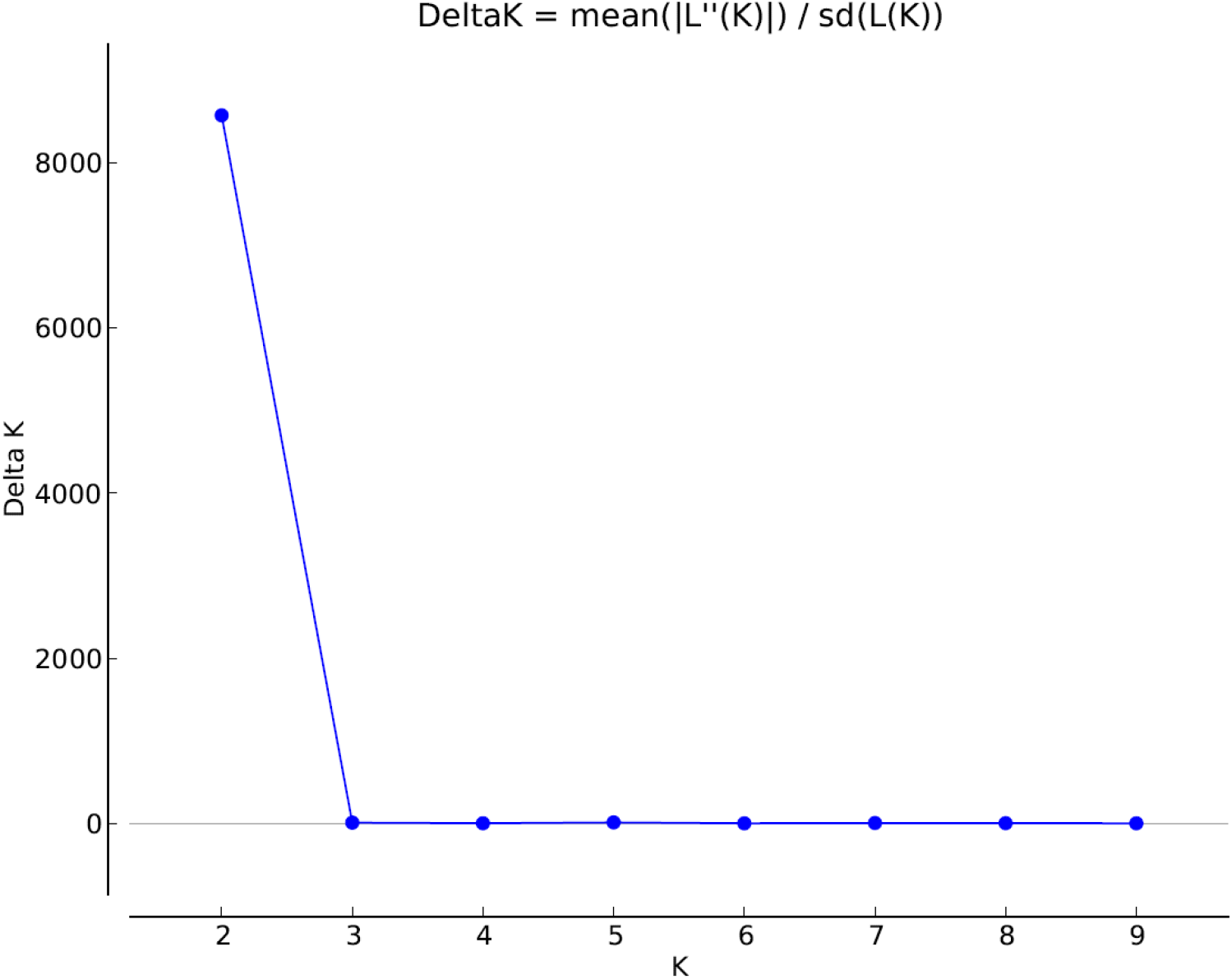
ΔK method plot with the optimal K of 2 for MPB, calculated in CLUMPAK using the Evanno method.

**Figure Error! No text of specified style in document..4:**
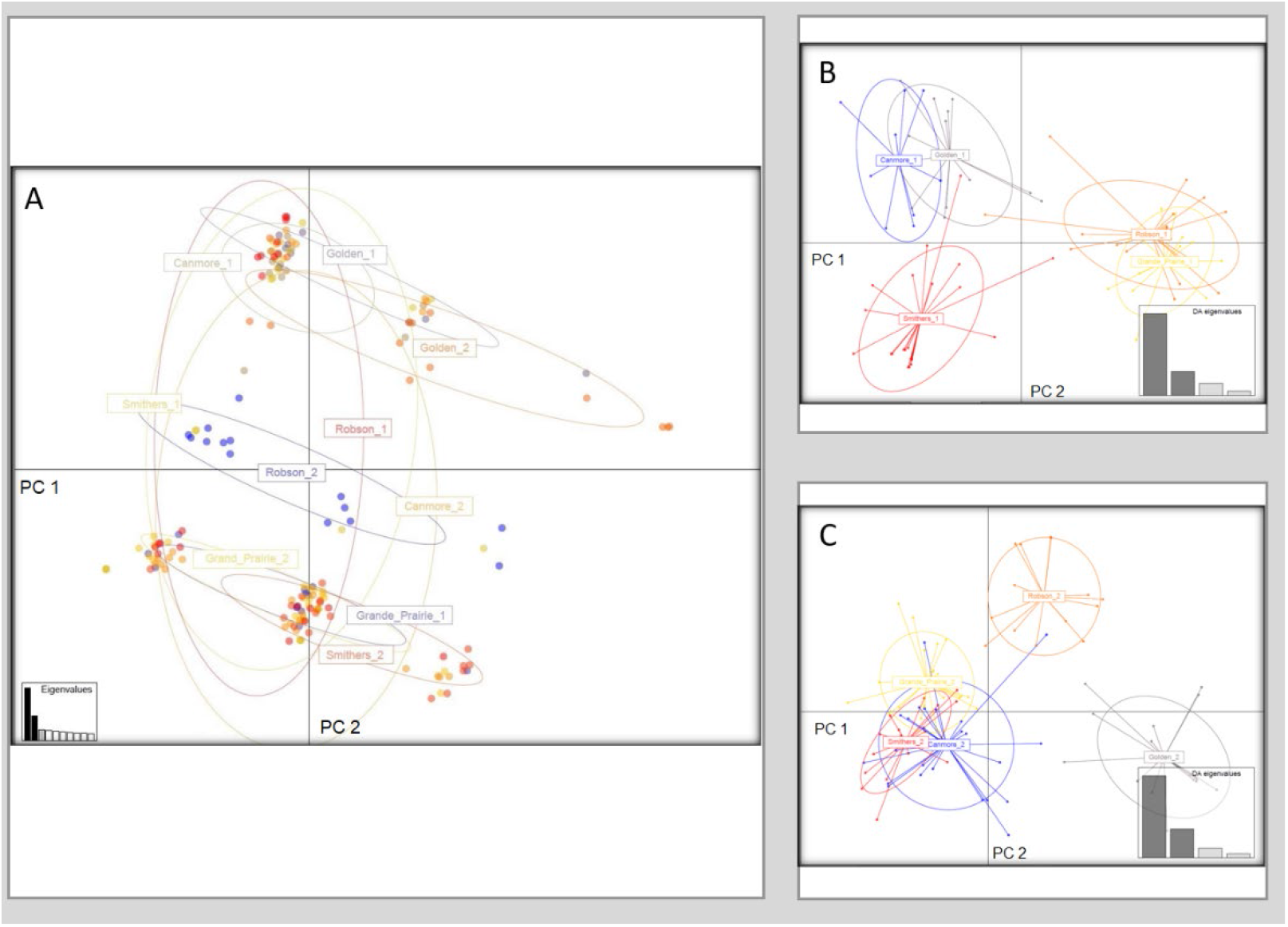
A) PCA of all MPB sites showing PC 1 and PC 2 and respective eigenvalues. B) DAPC scatterplot of MPB SNP genotypes displaying principal components 1 and 2 of years 2005/2007 C)DAPC scatterplot of MPB SNP genotypes displaying principal components 1 and 2 of year 2016.

**Figure Error! No text of specified style in document..5:**
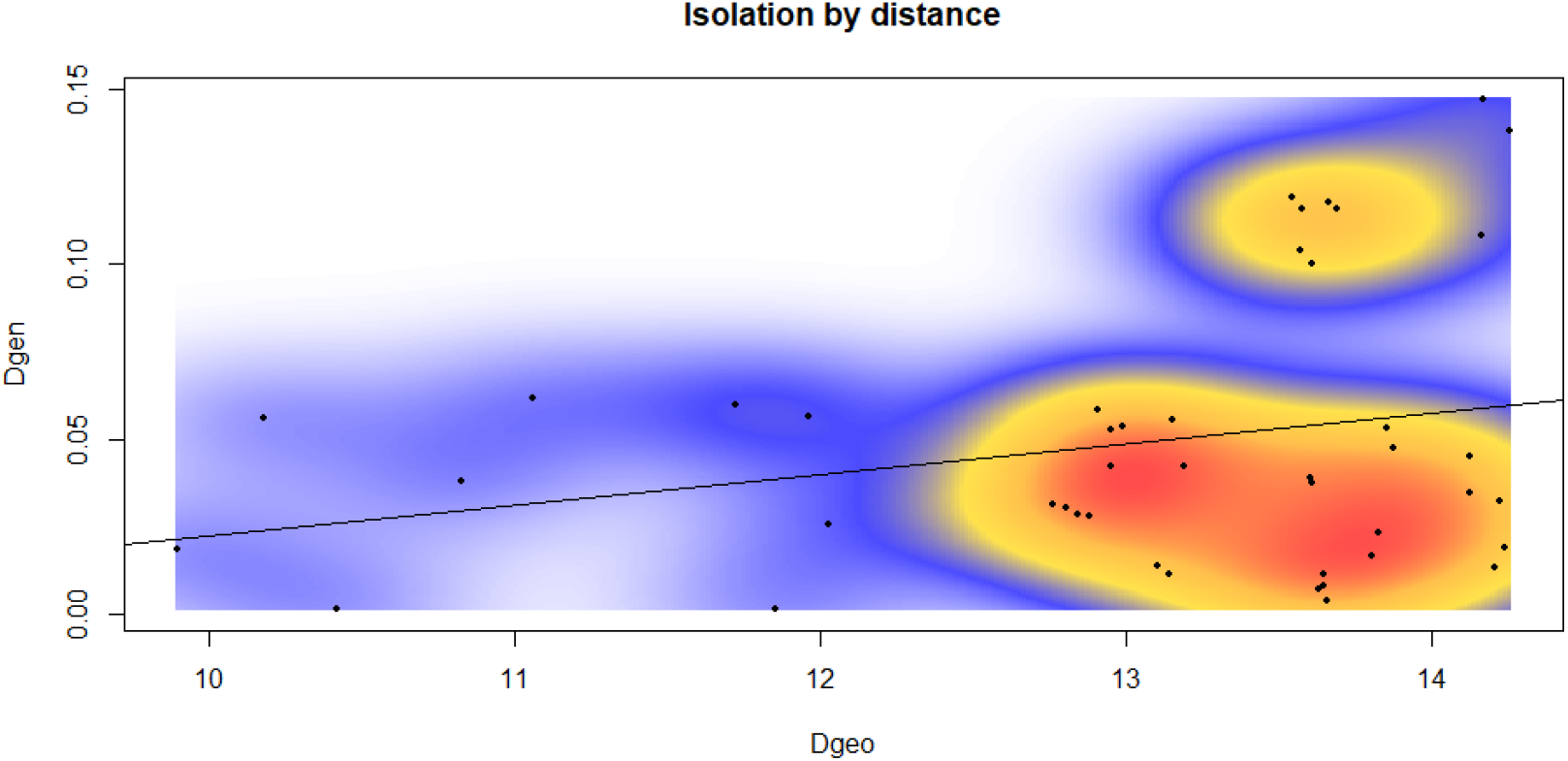
Isolation by distance for sites in 2005/2007 and 2016. The solid line through the points represents a Pearson-Moment correlation (non-significant) for the Mantel test (r2 = 0.2189, p = 0.102).

**Table Error! No text of specified style in document..1:**
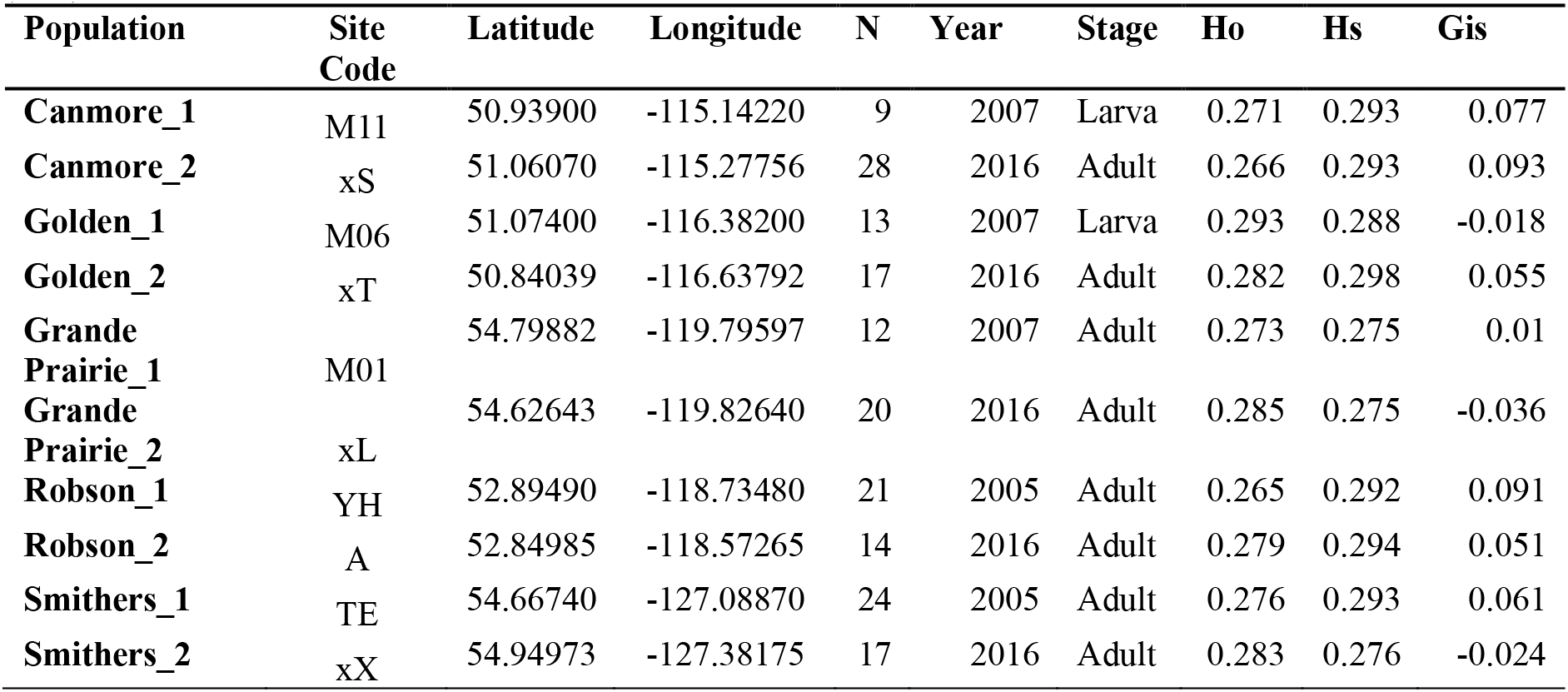
MPB site collection date, location, observed heterozygosity (Ho), heterozygosity within populations (Hs), and inbreeding coefficient (Gis).

**Table Error! No text of specified style in document..2:** Significance of pairwise Fst values based on P-values generated in Arlequin (+ = P < 0.05).

|  | Robson_1 | Grand Prairie_1 | Golden_1 | Canmore_1 | Smithers_1 | Smithers_2 | Robson_2 | Grand Prairie_2 | Golden_2 | Canmore_2 |
| --- | --- | --- | --- | --- | --- | --- | --- | --- | --- | --- |
| Robson_1 |  |  |  |  |  |  |  |  |  |  |
| Grand Prairie_1 | + |  |  |  |  |  |  |  |  |  |
| Golden_1 | + | + |  |  |  |  |  |  |  |  |
| Canmore_1 | + | + | - |  |  |  |  |  |  |  |
| Smithers_1 | + | + | + | + |  |  |  |  |  |  |
| Smithers_2 | + | + | + | + | + |  |  |  |  |  |
| Robson_2 | + | + | + | + | + | + |  |  |  |  |
| Grand Prairie_2 | + | - | + | + | + | + | + |  |  |  |
| Golden_2 | + | + | + | + | + | + | + | + |  |  |
| Canmore_2 | + | - | + | + | + | + | + | + | + |  |

**Table Error! No text of specified style in document..3:** Heatmap of Pairwise Fst values for MPB sites generated in Arlequin. Red indicates the highest levels of Fst while green indicates the lowest.

|  | Robson_<br>1 | Grand<br>Prairie 1 | Golden_<br>1 | Canmore_<br>1 | Smithers_<br>1 | Smithers_<br>2 | Robson_<br>2 | Grand<br>Prairie 2 | Golden_<br>2 | Canmore_<br>2 |
| --- | --- | --- | --- | --- | --- | --- | --- | --- | --- | --- |
| Robson_1 | 0.00 |  |  |  |  |  |  |  |  |  |
| Grand Prairie_1 | 0.03 | 0.00 |  |  |  |  |  |  |  |  |
| Golden_1 | 0.04 | 0.11 | 0.00 |  |  |  |  |  |  |  |
| Canmore_1 | 0.04 | 0.11 | 0.00 | 0.00 |  |  |  |  |  |  |
| Smithers_1 | 0.02 | 0.04 | 0.03 | 0.03 | 0.00 |  |  |  |  |  |
| Smithers_2 | 0.06 | 0.01 | 0.13 | 0.14 | 0.06 | 0.00 |  |  |  |  |
| Robson_2 | 0.01 | 0.02 | 0.05 | 0.05 | 0.02 | 0.04 | 0.00 |  |  |  |
| Grand Prairie_2 | 0.03 | 0.00 | 0.11 | 0.11 | 0.04 | 0.01 | 0.02 | 0.00 |  |  |
| Golden_2 | 0.05 | 0.09 | 0.03 | 0.04 | 0.04 | 0.10 | 0.05 | 0.09 | 0.00 |  |
| Canmore_2 | 0.01 | 0.00 | 0.06 | 0.06 | 0.02 | 0.02 | 0.01 | 0.01 | 0.05 | 0.00 |

**Table Error! No text of specified style in document..4:** Global AMOVA design for MPB individuals and results as a weighted average over loci (averaged over 4733 loci). An asterisk denotes significant values.

| Source of variation | SS | Var. comp. | % var | Fstat |
| --- | --- | --- | --- | --- |
| Among groups | 2305.948 | 3.44104 | 0.48023 | FCT :0.00480* |
| Among populations within groups | 12859.198 | 27.21377 | 3.79797 | FSC :0.03816* |
| Among individuals within populations | 113213.075 | 29.94989 | 4.17983 | FIS :0.04367* |
| Within individuals | 110537.500 | 655.92895 | 91.54196 | FIT :0.08458* |
| Total | 238915.720 | 716.53365 | --- |  |

## Acknowledgements

We would like to thank the British Columbia Ministry of Environment and Climate Change Strategy (Park Use Permit, Authorization # 107171), Alberta Agriculture and Forestry, and Robson Provincial Park staff for providing guidance on potential site locations. This research was supported in part by funding provided by the Natural Science and Engineering Research Council of Canada (grant no. NET GP 434810-12) to the TRIA Network. Computational work was supported by the cluster resources provided by Compute Canada (www.computecanada.ca).

